# Mobilisation of data from natural history collections can increase the quality and coverage of biodiversity information

**DOI:** 10.1101/2024.11.20.624468

**Authors:** Bryony Blades, Cristina Ronquillo, Joaquín Hortal

**Affiliations:** Department of Biology, University of Oxford, Oxford, United Kingdom; African Natural History Research Trust (ANHRT), Kingsland, United Kingdom; Department of Biogeography & Global Change, Museo Nacional de Ciencias Naturales (MNCN-CSIC), Madrid, Spain

**Keywords:** Biodiversity information, data coverage, digitisation, environmental bias, GBIF, mobilisation, museum, natural history collections, spatial bias

## Abstract

**Aim:** Quantify potential gains to insect data on the Global Biodiversity Information Facility (GBIF) through further digitisation of natural history collections, assess to what degree this would fill biases in spatial and environmental record coverage, and deepen understanding of environmental bias with regard to climate rarity.

**Location:** Afrotropical realm (mainland only)

**Time period:** 1814-2022

**Major taxa studied:** *Catharsius* Hope, 1837 (Coleoptera: Scarabaeidae)

**Methods:** We compared inventory completeness of Afrotropical *Catharsius* GBIF data to a dataset which combined these with records from a recent taxonomic revision. We analysed how this improved dataset reduced regional and environmental bias in the distribution of occurrence records using an approach that identifies well-surveyed spatial units of 100×100km as well as emerging techniques to classify rarity of climates.

**Results:** The number of cells for which inventory completeness could be calculated, as well as coverage of climate types by “well-sampled” cells, increased three-fold when using the combined set compared to the GBIF set. Improvements to sampling in Central and Western Africa were particularly striking. Coverage of rare climates was similarly improved, as not a single well-sampled cell from the GBIF data alone occurred in the rarest climate types. Inclusion of further records from natural history collections increased the total number of occurrences, but also filled persisting spatial and environmental data gaps on GBIF.

**Main conclusions:** These findings support existing literature that suggests data gaps on GBIF are still pervasive, especially for insects and in the tropics and, so, is not yet ready to serve as a standalone data source for all taxa. Biases in spatial coverage of records translate to uneven sampling of environmental conditions, hindering our ability to describe the full breadth of species’ niches, especially so in climates that occur infrequently. However, we show that natural history collections hold the necessary information to fill many of these gaps, and their further digitisation should be a priority.

## Introduction

In recent decades, the concurrent intensification of technological advancement and the coupled climate and biodiversity crisis has given rise to massive mobilisation of biodiversity ‘big data’ (Newbold, 2010; Wüest et al., 2020). This surge in data availability, apparent in the 3 billion records now available on the Global Biodiversity Information Facility (GBIF) – the world’s largest online biodiversity information network – has allowed researchers to ask questions on previously impossible scales in fields such as conservation, biodiversity informatics, macroecology, disease biology, and taxonomy (Heberling et al., 2021). Understanding of species’ distributions, and methods with which to model them, have particularly benefitted, in turn advancing knowledge of evolutionary processes and conservation management (Acevedo et al., 2016; Elith & Franklin, 2013; Guisan & Thuiller, 2005; Soberón & Peterson, 2004).

Despite the scale of these advancements, our understanding of species’ distributions is still incomplete at all spatial scales. Labelled the ‘Wallacean shortfall’ (Lomolino, 2004), knowledge gaps arise from uneven sampling effort and result in biases in spatial, temporal, climatic, and taxonomic coverage of records with locational information (Collen et al., 2008; Hortal et al., 2015; Oliver et al., 2021; Sporbert et al., 2019; Troudet et al., 2017). Despite their richness, data deficits are particularly pervasive in the tropics, especially so outside of protected areas, driven by a lack of appeal to surveyors, tough environmental conditions, limited accessibility, poor infrastructure or security, and a lack of local capacity such as academic institutions or funds (Amano & Sutherland, 2013; Araujo et al., 2022; Rocha-Ortega et al., 2021; Siddig, 2019; Yesson et al., 2007).

Deficits and spatial biases especially critical for invertebrates, with only 12% of the world’s territory sampled for insects (Rocha-Ortega et al., 2021), and a likely shortfall of over 200 million occurrences (Troudet et al., 2017), leading to particularly low inventory completeness – referring to how comprehensively biodiversity has been surveyed and recorded – even in scenarios where records are numerous (García-Rosello et al., 2023; Iannella et al., 2019; Sánchez-Fernández et al., 2021). Furthermore, record unevenness in space can also result in their biased distribution across environmental gradients, affecting our ability to characterise the real breadth of species’ fundamental niches (Hortal et al., 2008), although this is not always the case (Newbold, 2010). When GBIF insect and arachnid data were compared with alternate data sources such as taxonomic bibliographies and natural history collections, they were outperformed in the generation of ranges and climatic niches, and IUCN Red List classifications, despite the abundance of records (Beck et al., 2013; Shirey et al., 2019). Similarly, occurrences from systematic surveys outperformed GBIF data in the identification of well-surveyed cells for birds and fish in the United States, as they were more numerous and less spatially and environmentally biased (Troia & McManamay, 2016).

The way in which these same biases are reflected in the inventory completeness of insects has been investigated, but these studies generally compile exhaustive databases from a number of sources to quantify overall sampling effort as fully as possible (Ballesteros-Mejia et al., 2013; Romo et al., 2006; Sánchez-Fernández et al., 2008, 2022; Shirey et al., 2021). Much less is known about insect inventories derived only from GBIF data (see García-Rosello et al., 2023; Rocha-Ortega et al., 2021), and even less on how severely these are subject to spatial and environmental bias (but, see Girardello et al., 2019; Troia & McManamay, 2016). Additionally, whether inventory completeness is biased towards climate conditions that occur frequently is seldom analysed (but, see Ronquillo et al., 2020; Sobral-Souza et al., 2021), despite evidence that GBIF data is lacking in rare, locally-restricted taxa (Beck et al., 2013).

Given that species distribution modelling was the most prevalent topic in papers published using GBIF data between 2003 and 2019 (Heberling et al., 2021), and the way in which these correlative tools use climatic and species occurrence data to derive species–environment relationships, it is problematic that research to date has not yet fully determined how significantly the completeness of GBIF’s insect inventories is biased in climatic space. Accurate estimations of data completeness and coverage are essential in creating reliable models for species distributions and biodiversity patterns (Troia & McManamay, 2016, and references therein), and can be used to generate biogeographical maps of ignorance to improve predictions (Tessarolo et al., 2021). Without confidence in model accuracy, and an in-depth understanding of where they are least reliable, it is difficult to be sure that biased or erroneous results are not informing theoretical and practical applications in ecology and conservation.

Although intensified digitisation and mobilisation (meaning digitised and made widely available through an online database) of natural history collections onto GBIF has begun to help fill data gaps, especially with contributions from smaller herbaria and institutions, as well as private collections (Araujo et al., 2022; Beck et al., 2013; Yesson et al., 2007), much remains to be done (Hardy et al., 2023; Popov et al., 2021). Fortunately, data compilation for purposes such as taxonomic revisions often utilises collections that have heretofore not been mobilised. By quantifying how much inventories are improved when such collections are included, a better understanding of the remaining deficits in GBIF data can be reached.

In this study, spatial and environmental bias in GBIF insect inventories is evaluated using data on dung beetles, which are known to act as a proxy for general biodiversity (Spector, 2006). We determine how significantly bias could be reduced through the integration into GBIF of a dataset independently compiled for a taxonomic revision of the Afrotropical members of the genus *Catharsius* Hope, 1837 (Coleoptera: Scarabaeidae) (BLINDED, 2024). In particular, we investigate inventory completeness and to what extent this is affected by climatic conditions and rare climates.

## Materials and Methods

### Study area and taxon

The study area is the African mainland of the Afrotropical biogeographical realm spanning 17.5° W – 51° E, 21° N – 35° S. The study region exhibits a wide breath of climatic conditions, including eight biomes and 91 ecoregions, encompassing tropical forest to xeric shrublands. Using QGIS version 3.22.9-Białowieża (2021), a shapefile of the realm (Olson et al., 2001; World Wildlife Fund, 2012) was modified to include only the mainland of Africa, categorising countries by region using the UN M49 standard (United Nations Statistics Division, n.d.) as a reference (see Appendix S1 in Supporting Information). This shapefile was then used to create a grid of 100km x 100km cells for the statistical analyses.

*Catharsius* is a genus of large copro- and necrophagous dung beetles with species distributed across both Africa and Asia. The Afrotropical members are currently being revised in the largest ever revision of any group of dung beetles in the world, for which an extensive collection of distributional data has been compiled from natural history collections (H. Takano, African Natural History Research Trust, personal communication; BLINDED, 2024), and it is these records that are described below.

### Occurrence data

To demonstrate the potential value of integrating unmobilised natural history collection data to GBIF, two datasets were used. Firstly, all occurrences for *Catharsius* were downloaded from GBIF (13/11/2023) and subjected to a pre-processing procedure to ensure they did not fall foul of known GBIF data quality issues, as follows. Occurrences were filtered taxonomically to include only those identified to species, and geographically to include only those with precise and accurate coordinate information that did not suggest specimens were located in biodiversity institutions, the sea, in the centroid of a country, or in countries other than those listed on the physical label. These steps help to ensure that metadata is robust and corresponds to the right species (Ronquillo et al., 2024), and further specifics can be found in Appendix S2. The taxonomy of the resulting occurrences was standardised according to the most recent revision (BLINDED, 2024; see Appendix S2) and cropped to the same extent as the bioclimatic variables, and this is henceforth referred to as the ‘GBIF set’.

Secondly, a combined dataset was created by joining the GBIF set and a set of species names, coordinates, years, and counts that had previously been extracted from the text of the taxonomic revision- the ‘revision set’ (BLINDED, 2024). As part of the extraction process, all occurrences had been manually inspected to ensure they fell on land and in countries that were expected for that species. A small number of errors from typing inaccuracies were corrected with the expert help of the original taxonomist. These records were cropped to the study extent before merging with GBIF data and duplicate entries manually removed (see Appendix S2). This is henceforth referred to as the ‘combined set’.

### Statistical analyses

To assess survey completeness in all grid cells of the study area, the package ‘KnowBR’ version 2.2 (Guisande & Lobo, 2023) was used. By generating species accumulation curves – plots representing the cumulative number of species observed as a function of the cumulative number of samples collected – it compares the number of observed species to the number of predicted species per spatial unit (here, 100 km x 100km cells). This determines a percentage of inventory completeness, i.e. how many of the species that are likely to be present in each cell have been recorded as such (Lobo et al., 2018). Analysis was carried out separately for both the GBIF and combined sets, and then cells with >20 records, a completeness score > 75%, and a ratio of occurrences to species >5 were identified as ‘well-sampled’ (WS) in each set. The regional shapefile was used to quantify bias of inventory completeness and WS cells for both sets in terms of how evenly they were distributed in each region.

As unevenness in the spatial coverage of occurrence data has been shown to result in biased sampling of environmental conditions (Hortal et al., 2008), the WS cells for each set were then used to evaluate if comprehensive sampling has been conducted across the full spectrum of potential environmental conditions in the study region. To describe environmental variations in the study area, we used the 19 bioclimatic variables from WorldClim, which represent trends, seasonality, and limiting environmental factors of temperature and precipitation derived from aggregating monthly data from the period 1970–2000 (Fick & Hijmans, 2017; WorldClim, 2020; see Appendix S3). These were downloaded at a resolution of 30 seconds (approximately 1km at the equator), and aggregated to 0.83 degrees (approximately 100km at the equator), the resolution of the grid used for spatial coverage, and cropped to the study extent.

A principal components analysis (PCA) of the study area was carried out in R package ‘psych’ version 2.4.1 (Revelle, 2024) to reduce the dimensionality of the climate data to two axes (PC1 and PC2) that captured 73% of the variation. PC1 predominantly characterised levels of precipitation and variability in temperature, displaying a gradient from the high rainfall and more stable temperatures of tropical forests, to the drier desert areas with greater annual and daily temperature ranges. PC2 broadly described a temperature gradient from the warmer north and low elevations, to the cooler south and high elevations, as well as high to low precipitation seasonality to a lesser degree. By converting PCA values into classes, 33 distinct climate types were identified in the study area (see Appendix S3).

Following methods to measure climatic coverage of sampling in Sobral-Souza (2021) and Ronquillo et al. (2020), the classes – or ‘climate types’ – and the frequency at which each was observed in the study region was quantified. The climate types that the WS cells fell in were identified and the niche overlap between conditions in these locations and the study region as a whole was calculated using Schoener’s *D*. This returns a value of between 0, for no overlap, and 1, for a complete overlap (Schoener, 1970). Here, as WS cells are found in an increasing number of climate conditions, Schoener’s *D* would be expected to rise, indicating good sampling over a more comprehensive coverage of potential environmental conditions in the Afrotropical realm. Significance of *D* values was tested by comparing them against a null distribution, generated by randomly sampling 1000 sets of five (for the GBIF set) or 23 (for the combined set) occurrences (i.e. the same number of occurrences as WS cells for each set) and calculating the niche overlap of each sample with the study area.

To determine whether environmental conditions found in WS cells were an unbiased subset of those described by each PCA axis – i.e. whether better sampling is correlated with particular conditions – two sample Kolmogorov-Smirnov tests were run using the R base package ‘stats’. Used to evaluate whether two groups are sampled from the same distribution (Massey, 1951), the null hypothesis will be rejected if the WS cells sample certain climatic conditions at a rate that does not reflect their overall frequency in the study area. For example, if WS cells are found to disproportionately favour cooler climates, despite the presence of warmer climates in the realm.

Finally, min-max scaling was used to evaluate whether WS cells were found in climates that occur infrequently in the study region, or are ‘rare’. For this, a rarity value between 0 (common) and 1 (rare) was assigned to each climate type, depending on its relative frequency in the study region. The rate at which each type was sampled by WS cells was then compared to its overall density in the study region, as such determining whether good sampling was biased towards common or rare climates.

All analyses were conducted in R version 4.2.1 (R Core Team, 2022), using RStudio version 2023.12.1.402 (Posit Team, 2024), and code adapted from Ronquillo (2023).

## Results

### Pre-processing of occurrence data

Downloading all records for *Catharsius* from GBIF returned a dataset of 4270 entries (GBIF, 13/11/2023), pertaining to 72 species; more results of the quality assurance filtering and taxonomic standardisation procedures can be found in Appendix S2. After pre-processing, the GBIF set totalled 1686 entries, corresponding to 3915 specimens belonging to 50 species. The revision dataset totalled 4979 entries, corresponding to 15,943 specimens belonging to 146 species. Upon combination with the GBIF dataset, 489 duplicate entries were removed from the former, resulting in a combined dataset of 6174 entries, corresponding to 18,043 specimens belonging to 146 species. As such, the revision set contributed the overwhelming majority of information to the combined set with GBIF having contributed only 27.3% of entries, 21.7% of specimens, and no new species.

### Statistical analyses

A total of 94 cells, out of a potential 1867 (5%), contained sufficient information to compute inventory completeness for the GBIF set, which are predominantly concentrated in the northeast of South Africa (Figure 1). Regionally, 51 of these cells are found in Southern Africa, 41 in Eastern Africa, two in Central Africa, and none in Western Africa. Contrastingly, inventory completeness could be computed for 314 (16.82%) with the combined set, an increase of 220. Of these, 75 are found in Southern Africa, 128 in Eastern Africa, 76 in Central Africa, and 35 in Western Africa, illustrating a reduced regional sampling bias. Some cells for the combined set lay across the border between two regions, in which case the centre point of the grid square was the decider.

**Figure 1:**
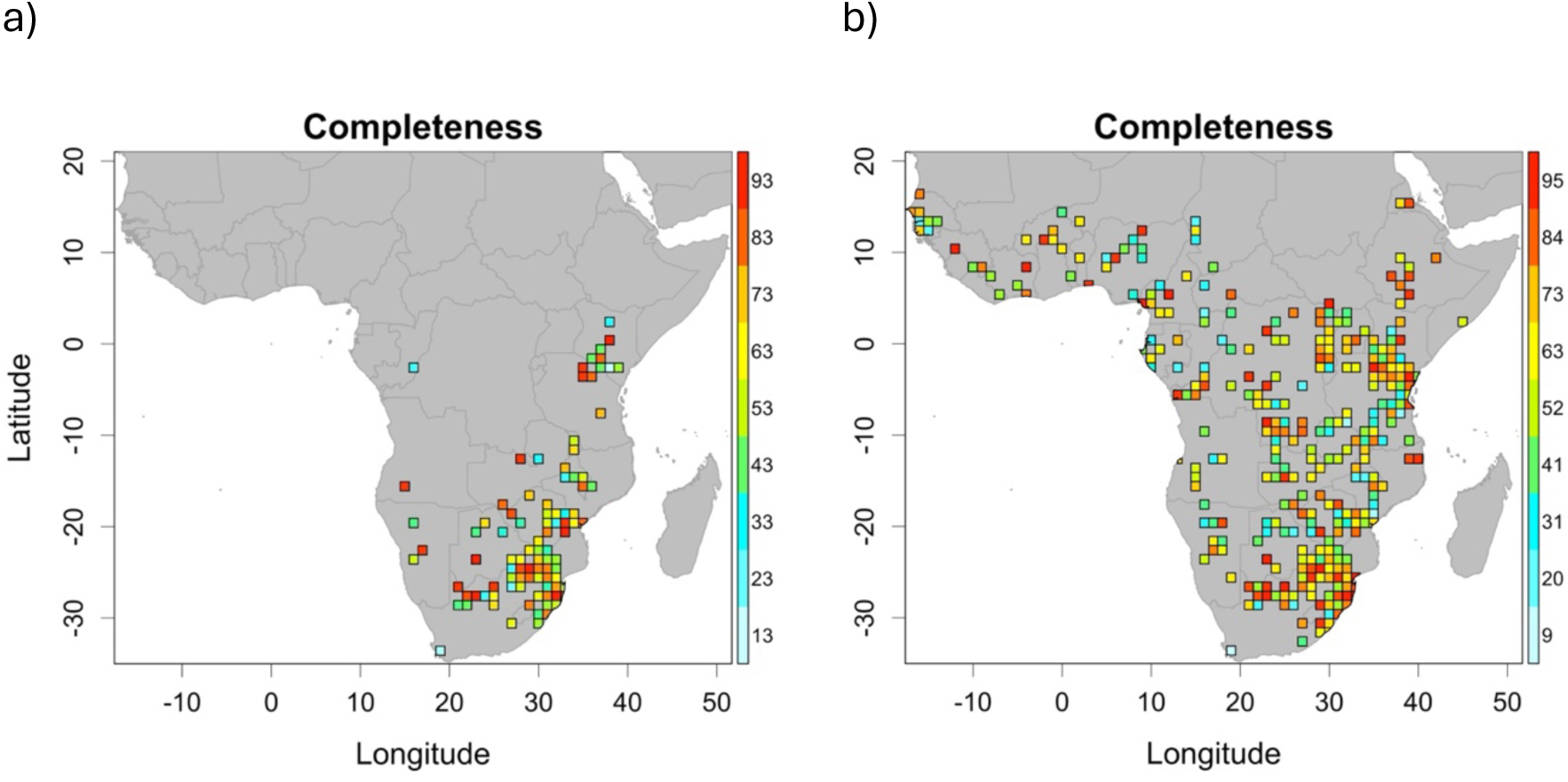
Completeness of *Catharsius* inventories in the mainland Afrotropical realm using the a) GBIF set and b) combined set. Coloured cells are those for which data were sufficient to calculate completeness, with warmer colours indicating higher completeness percentage

Cells with high completeness values were also concentrated in South Africa, with some further representation in Southern and Eastern Africa for the GBIF set, but a cross-realm spread for the combined set. The single grid cell in Central Africa with a high completeness value in the GBIF set was no longer considered to be so well-completed in the context of a more extensive dataset.

Only five cells fulfilled the criteria to be considered well-sampled using the GBIF set, with four in the northeast of South Africa, and one in Kenya; just 0.27% of 1867 potential cells, and 2 countries. Using the combined set, twenty-three WS cells (1.23%) were identified across 11 countries (8 in South Africa, 3 in the Democratic Republic of Congo, 3 in Mozambique, 2 in Kenya, and 1 in each of Senegal, Cote d’Ivoire, Cameroon, South Sudan, Zimbabwe, Uganda, and Tanzania). Although South Africa is also comparatively overrepresented, and 7 out of these 11 countries only returned a single WS cell, the combined set generated 4.5 times more WS cells in 5.5 times more countries than the GBIF set.

Only four out of a potential 33 distinct climate types (see Appendix S3) were found in WS cells from the GBIF set (12.12%), as opposed to 12 for the combined set (36.36%). The niche overlap (*D*) between WS cells and the study region as a whole was 0.238 (*p* = 1) and 0.468 (*p* = 1) for the GBIF and combined set, respectively.

Kolmogorov-Smirnov tests show WS cells for both sets as unbiased in PCA1 (GBIF: *D* = 0.30, p = 0.75; combined: *D* = 0.21, p = 0.26) but biased in PCA2 (GBIF: *D* = 0.63, p = 0.04; combined: *D* = 0.35, p = 0.0083). As such, whilst both are representative samples of potential levels of precipitation and temperature variability, they oversample moderate and cooler temperatures, with comparatively little representation of warmer climates in the north (Figure 3).

No WS cells from the GBIF set were found in the rarest of climates (rarity > 0.8), and both very common and moderately common climate types were overrepresented compared to their density in the study region. Whilst WS cells from the combined set also under-sampled the rarest of climates, it was to a lesser degree, and common and moderately common climate types were sampled at a rate more representative of their overall density in the study region (Figure 4). Improved sampling in Central and Eastern Africa was responsible for coverage of the rarest climates by combined set WS cells, and the failure of GBIF WS cells to sample any of these was despite their both being most prevalent in South Africa (Figure 5).

## Discussion

This study provides novel insights into inventory completeness of the world’s biggest online biodiversity data network. By quantifying coverage improvement achieved with the addition of further natural history collection records, this comparative approach assesses not just how much value is still missing for insects on GBIF, but also the scale of potential gains from further record digitisation. In particular, it has shown that GBIF occurrence records are biased towards the southern and eastern areas of the Afrotropical realm and, consequently, fail to sample across the full range of potential environmental conditions. This leads to poor coverage of warmer climates, and rare climate types. Whilst inclusion of further natural history collection data does not entirely remove sampling bias, it greatly reduces unevenness in spatial and environmental sampling.

Inventory completeness for GBIF insect data is well-documented as being poor worldwide, particularly so outside of Europe (García-Rosello et al., 2023; Rocha-Ortega et al., 2021), and this is reflected very clearly in these results. Here, GBIF data is only sufficient to compute completeness for 5% of grid cells and, even then, many of these return poor values and are strongly regionally biased, the pattern of which is broadly comparable with general inventory completeness of insects on GBIF (Figure 2c, García-Rosello et al., 2023 p. 493) . The combined set, though, allows completeness to be computed for over three times as many cells, and regional gaps are filled. It provides enough data to compute values in 74 and 35 more cells in Central and Western Africa, respectively, than the GBIF set, and also generated highly completed cells more evenly across the realm.

**Figure 2:**
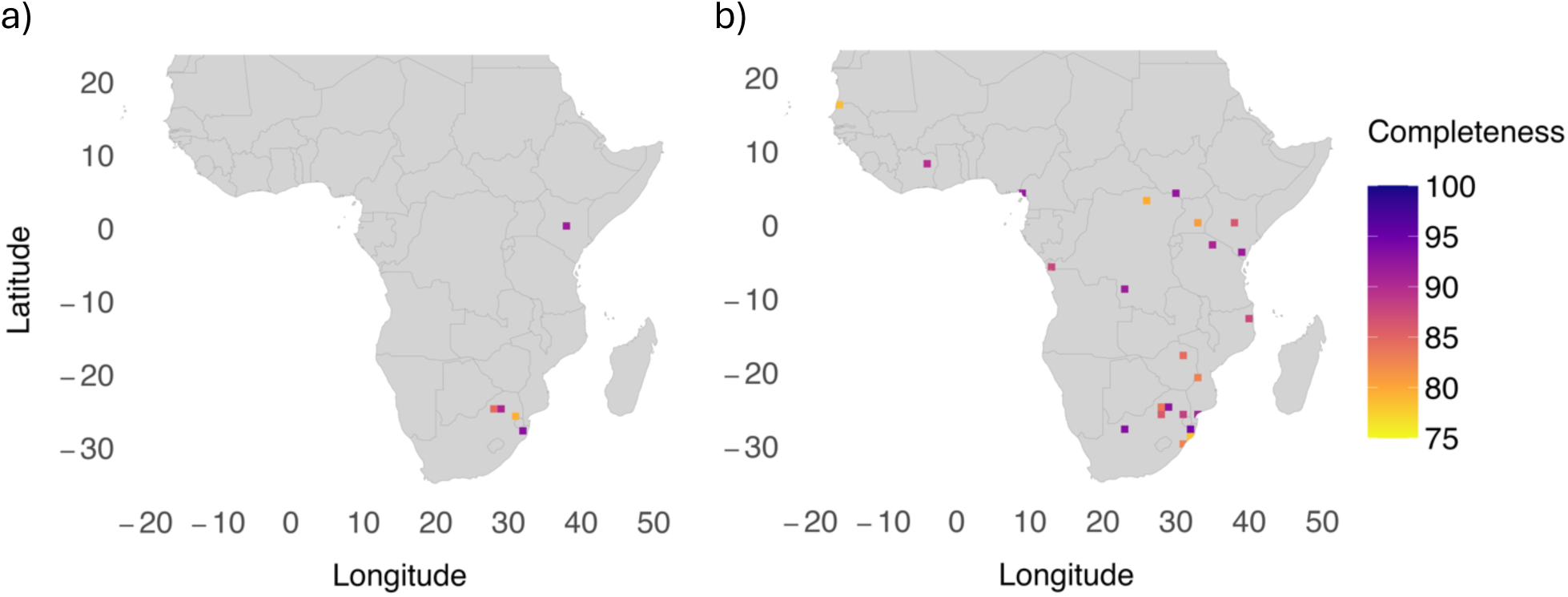
Cells in the mainland Afrotropical realm in which *Catharsius* is well-sampled using a) the GBIF set and b) the combined set, where well-sampled is defined as containing >20 records, completeness >75%, and a ratio of occurrences to species >5. Colour indicates completeness percentage, with darker colours identifying higher completeness

**Figure 3:**
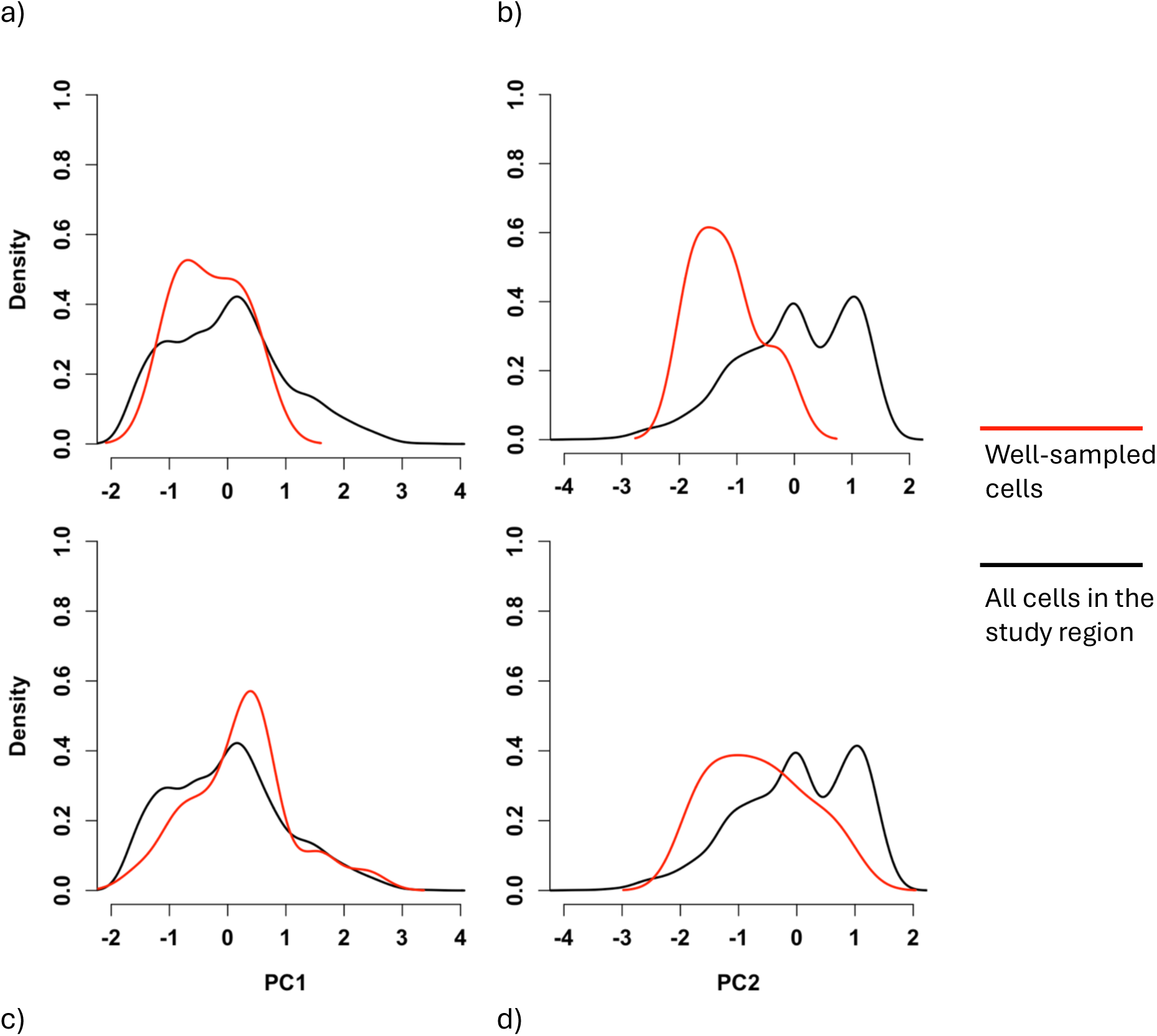
Smoothed kernel density estimate of the distribution of principal component axes (PC1 and PC2) scores for the GBIF set (a and b) and the combined set (c and d), with the y-axis representing relative density. The red and black lines illustrate the continuous distribution of PCA values found in the *Catharsius* well-sampled cells and all cells in the mainland Afrotropical realm, respectively.

**Figure 4:**
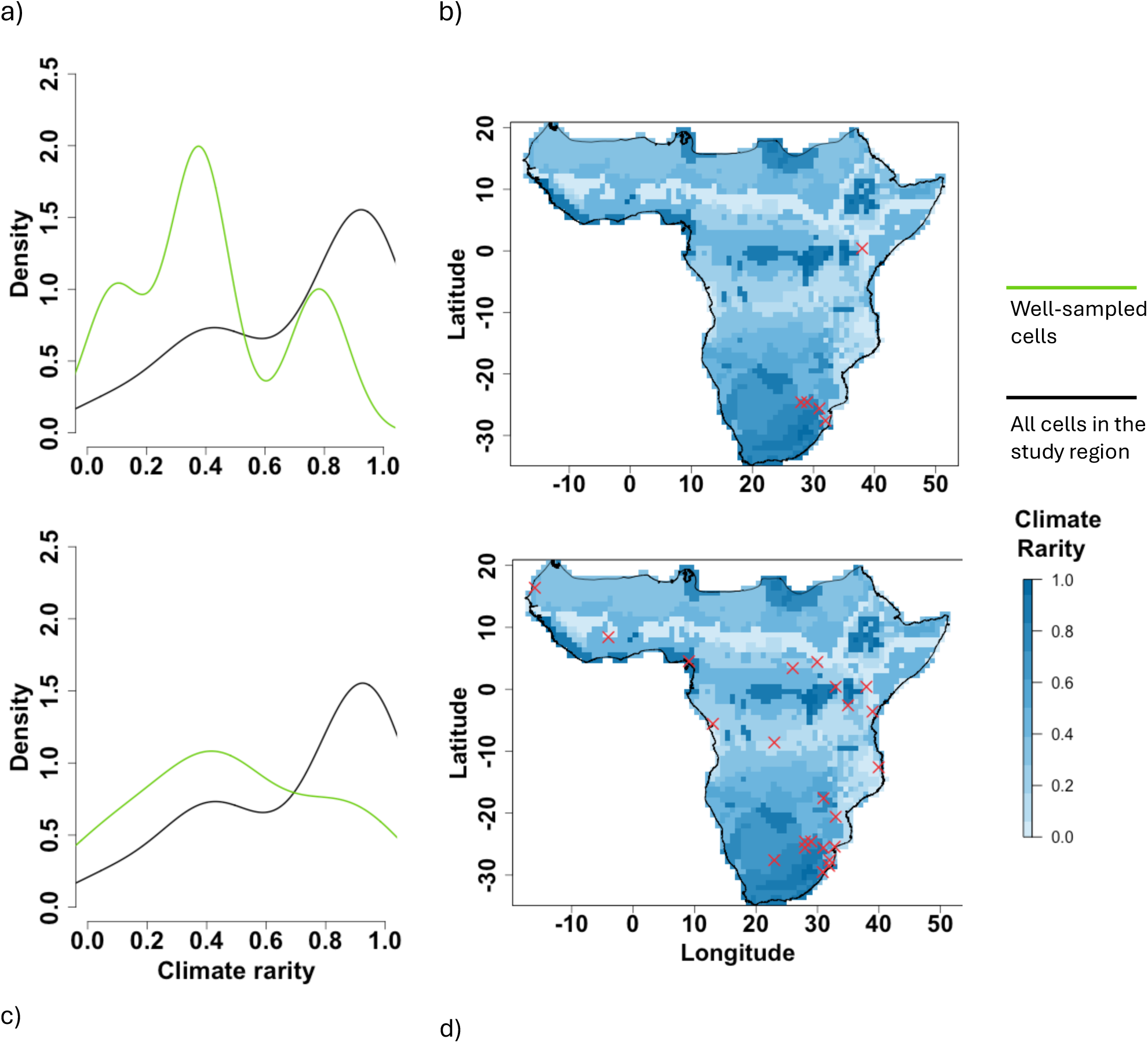
Smoothed kernel density estimate of the distribution of climate rarity scores, between 0 (common) and rare (1), for the GBIF set (a) and the combined set (c), with the y-axis representing relative density. The green and black lines illustrate the continuous distribution of climate rarity values found in the *Catharsius* well-sampled cells and all cells in the mainland Afrotropical realm, respectively. Maps illustrate the geographic distribution of climate rarity, with dark colours representing rarer climates, and well-sampled *Catharsius* cells for the GBIF set (b) and the combined set (d).

Notably, the two highly completed GBIF cells in Central Africa intersect Mupa and Bicuari National Parks in Angola, which is consistent with Girardello *et al*. (2019) who found gaps in GBIF butterfly inventories to correlate with low density of protected areas. In fact, these results support that geographic biases in GBIF insect inventory completeness are comparable to those in GBIF raw data (García-Rosello et al., 2023), as they identify deficiencies driven by survey area attractiveness and socioeconomic factors. The GBIF set generated 4.5 times less WS cells than the combined set, and their exclusive location in South Africa is consistent with findings that GBIF sampling coverage is strongly related to GDP per capita (Amano & Sutherland, 2013; Hughes et al., 2021; IMF, 2024), and shows similarities with the work of Stropp et al. (2016) on seed plants, evidencing that many biases in biodiversity knowledge are pervasive across groups. Contrastingly, almost a third of combined set WS cells were generated in countries ranked in the bottom 10 for 2024 GDP per capita (Democratic Republic of Congo, Mozambique, and South Sudan) (IMF, 2024). Species data used in this study are not from 2024, but this does pose the question of whether bias in GBIF data driven by economic inequality could be reduced by further digitisation of natural history collections, and this should be specifically investigated.

In light of the stark regional biases in GBIF WS cells, it is not surprising that coverage of climatic conditions also fall short. Well-sampled cells from the combined set covered more climate types of, and had a higher niche overlap with, the overall study region, demonstrating the potential value added to ecological inferences through increased record digitisation. That said, non-significant D values for both the GBIF set and the combined set indicate that WS cells are found in common climate types for both, despite general coverage improvement through integration of further data. These results align with studies in which systematic surveys and natural history collections outperformed GBIF insect data in terms of survey coverage but also derivation of ranges and climatic niches (Beck et al., 2013; Troia & McManamay, 2016). It is concerning that the scale of bias in GBIF insect data is such that it compromises its usefulness despite the sheer number of records now accessible. This has also been observed in other taxa (Araujo et al., 2022; Shirey et al., 2019) and, whilst its severity is likely not consistent across groups, it is probable that GBIF is not yet comprehensive enough to be used as an exclusive data source in modelling biodiversity patterns, and should instead be used as a resource within a wider suite. Others have also called for the integration of biodiversity databases in one place to this same end (Araujo et al., 2022).

A particular weakness of GBIF here was that, whilst most climate types in this study occurred infrequently, this was not reflected in sampling; the GBIF WS cells did not occur in any of the rarest climate types, despite both being most prevalent in South Africa. This risks missing species that may be specialised to infrequent climatic conditions and, in the case of such habitat specificity, at risk on multiple fronts: smaller geographic range, few populations, and specialised niche conditions (Işik, 2011). This problematic given the commonness of climate rarity in the Afrotropical realm, and means that data on many such rare species may not be made accessible before a conservation plan can be designed to combat their decline, greatly damaging functional diversity (Leitão et al., 2016). Furthermore, prioritising rare climates has recently been shown to result in higher species richness in protected area planning (Kim & Choe, 2024), and so it follows that efforts to improve data coverage in these areas should be paramount. Half of GBIF’s observations are from citizen science (Beck et al., 2013; Waller, 2019), which is known to under-sample invertebrates and suffer from further spatio-taxonomic biases (Chandler et al., 2017; Fontaine et al., 2022; Shirey et al., 2021; Theobald et al., 2015). On the other hand, small institutions and private collections contain distinctive information on species distributions, including for poorly sampled taxa and regions, such as the tropics, and the importance of digitisation and mobilisation of these data has been well-argued elsewhere (Araujo et al., 2022; Beck et al., 2013; Glon et al., 2017; Yesson et al., 2007). This study shows that the capacity to improve data coverage in rare climates is further reason to do so. Further, it pinpoints that a worthwhile direction for future research is a breakdown of data provenance, not just for GBIF but also other databases, to unpick the relative contributions of various biodiversity data sources in combatting knowledge shortfalls. This has been completed for Norway (Petersen et al., 2021), but expanding this methodology to data poor regions would be valuable.

Whilst the results in this study underline the importance of further natural history collection digitisation to ecology and conservation, removing data access obstacles will also benefit research areas such as invasive species management, medicine discovery, agricultural research and development, and mineral discovery, as well as provide knock-on benefits in education, public engagement, and informing data-driven policy (Hedrick et al., 2020; Popov et al., 2021; V. S. Smith et al., 2022). Lack of funding and resources are acknowledged as barriers to widescale digitisation (V. S. Smith et al., 2022), and the scale of the task is such that meaningful progress necessitates effective prioritisation. Methods to achieve this attribute varying weights to administrative criteria such as relevance, data quality, cost, and feasibility (Ahl et al., 2023), or more ecology-specific requirements to improve descriptions of biodiversity, including derivations of historical baselines to better understand anthropogenic change (Hedrick et al., 2020). Particularly pertinent is the recommendation to prioritise digitisation of specimens that will fill existing data gaps (Hedrick et al., 2020), such as the biases in centralised aggregators like GBIF, shown here.

Usefulness of digitised records, though, hinges on the quality of the data that is generated. Here, over half of the original GBIF data were removed during pre-processing as it did not meet the necessary standards for analysis (see Appendix S2), and concerns with geospatial errors, insufficient metadata, and taxonomic inaccuracies are well-documented (Ferro & Flick, 2015; Prudic et al., 2023; Rocha-Ortega et al., 2021; Ronquillo et al., 2020; Yesson et al., 2007). Misidentifications and poor taxonomy are particularly difficult to pinpoint and correct after mobilisation (Soberón et al., 2002), and this is especially so for little-known or cryptic invertebrate species, such as those in *Catharsius*. The revision set data was extracted from a rigorous, multi-year project to revise *Catharsius*’ taxonomy, so there were very few georeferencing errors and no taxonomic errors, according to this most recent understanding of the genus. The time-consuming nature of this work and declining taxonomic expertise (Hopkins & Freckleton, 2002; Hutchings, 2021; ‘Importance of Taxonomy’, 1946; Lagomarsino & Frost, 2020; Wägele et al., 2011), precludes this as an option for many studies, especially those that encompass thousands, or even millions, of occurrence records.

Methods to reduce GBIF taxonomic misidentifications have been tested with some success (B. E. Smith et al., 2016), but these still have distinct time and data demands of their own. Other tools to validate not just the taxonomic, but also geographic, temporal, and metadata accuracy of records are also promising (Ronquillo et al., 2024), but the quality of GBIF data is such that these processes often greatly reduce the number of usable data points, as seen in this study. It seems clear that the best way to avoid compounding existing flaws in data quality is with meticulous digitisation in the first place, including the allocation of resources to taxonomic verification or revision as part of the data preparation process.

The importance of insects to global biodiversity and ecosystem services cannot be understated (Noriega et al., 2018; Wagner et al., 2021), and that an unknown number may have already gone extinct (García-Rosello et al., 2023) paints a concerning picture for biodiversity in the Anthropocene. Efforts to understand the limitations of our knowledge are paramount. Whilst online mobilisation of biodiversity data on a massive scale has facilitated advancement in a number of fields (Heberling et al., 2021; Soberón & Peterson, 2004), and the rate of record digitisation has increased, gaps in coverage persist, and are particularly strong in the tropics and for invertebrates (Collen et al., 2008; Oliver et al., 2021; Sporbert et al., 2019; Troudet et al., 2017). This inherently affects the reliability of modelling predictions, potentially misinforming applications in ecology and conservation (Troia & McManamay, 2016). Research to date has found that completeness of insect inventories compiled from multiple sources varies across space and environment (Ballesteros-Mejia et al., 2013; Romo et al., 2006; Sánchez-Fernández et al., 2008, 2022; Shirey et al., 2021), but much less is understood about this using GBIF data alone (Girardello et al., 2019; Troia & McManamay, 2016). Research that seeks to better understand these flaws, and suggest solutions, not only improves theoretical knowledge of species distributions, but can potentially better inform practical applications, such as directing field work to areas where more data will maximise biogeographical inference. This study underlines that much value still stands to be gained by further digitisation of natural history collections, emphasising that GBIF is not yet ready to function as a standalone data source, but care must be taken to not compound existing data quality issues.

## Acknowledgements

We are particularly thankful to Hitoshi Takano for his unselfish release of unpublished data, and his guidance with taxonomic disambiguation of the classification of *Catharsius* dung beetles, without which this work wouldn’t have been possible. The authors also thank Tim Coulson, Elizabeth Jeffers, Michael Bonsall, and Robert Whittaker who commented on versions of this manuscript, and Richard Smith for his donation to the African Natural History Research Trust Scholarship.

## Data availability statement

Data and code supporting this paper are available on figshare and have a reserved DOI. For anonymity during peer-review, the data are cited in the manuscript as (BLINDED, 2024), and the following private link can be used to access them: https://figshare.com/s/a5fe8cf5587d2f06d87c

## Biosketches

**Bryony Blades** is a biogeography researcher working on describing distributions of tropical insects through species distribution models and landscape genomics, currently with a focus on Scarabaeinae in South-Central Africa. She also investigates how the quality and coverage of existing biodiversity data can be improved, specifically analysing the relationship between taxonomy, museum data, and modelling.

**Cristina Ronquillo** is an ecologist interested in macroecology, ecoinformatics, Geographic Information Systems and biodiversity databases. Her research interests include the assessment of data quality and information gaps and how it affects our biodiversity knowledge.

**Joaquín Hortal** is a biogeographer and community ecologist, working on understanding the effects of current and past environmental conditions and ecological processes on the geographic patterns of biodiversity, the role of species niche and ecological interactions in the organization of biological communities in time and space, and the use and misuse of biodiversity big data.

## Appendices – found in Supporting Information

S1 – Study Region

S2 – Occurrence data

S3 – Environmental data

