## Supplementary Information for "Mobilisation of data from natural history collections can increase the quality and coverage of biodiversity information"

### Appendix S1 – Study region

Figure S1.1: The study region

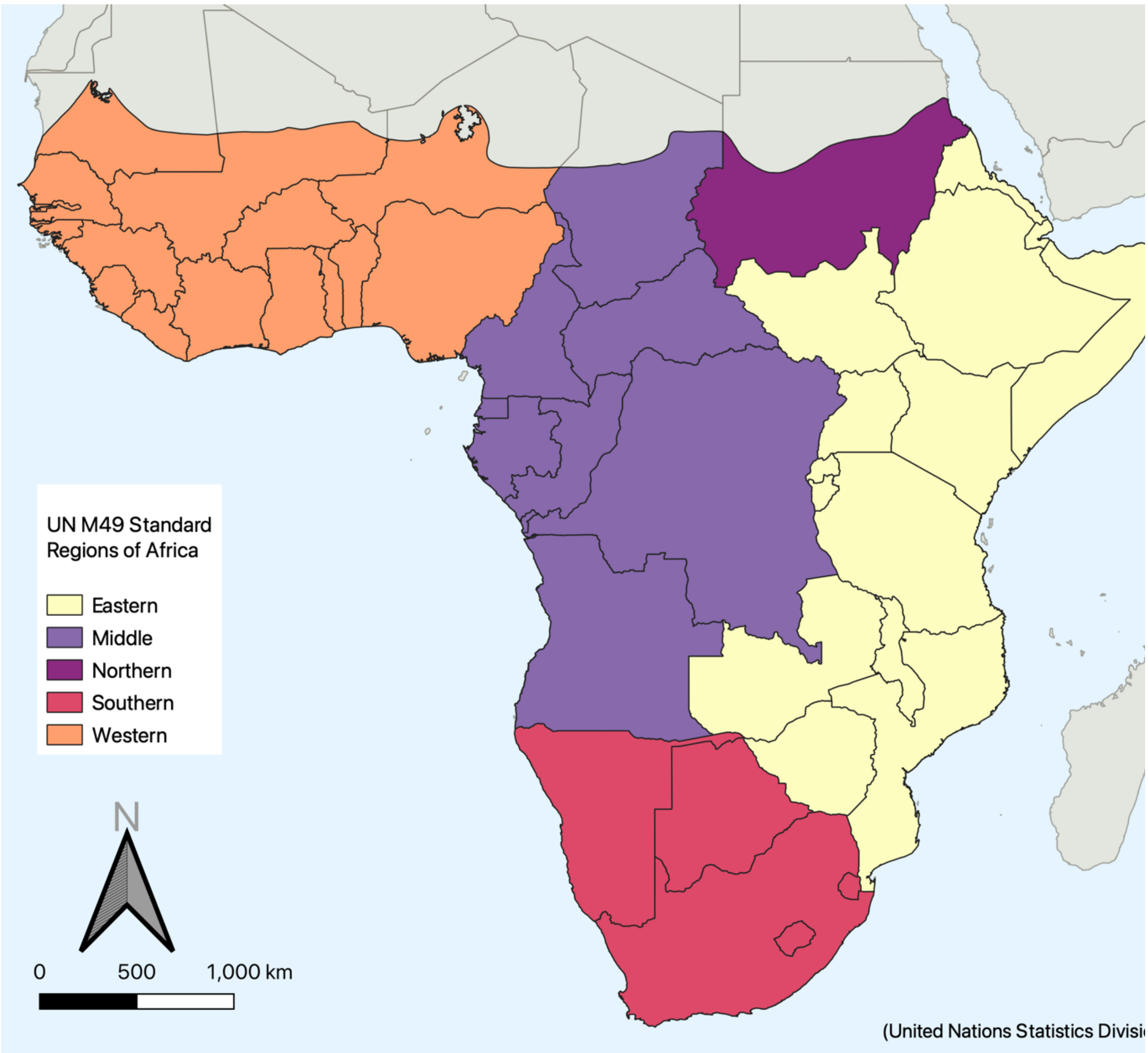

Figure S1.1: The mainland Afrotropical realm, created from World Wildlife Fund (2012), showing regions of Africa as designated by the UN M49 Standard (United Nations Statistics Division, n.d.). For the purposes of the study, the manuscript uses the term "Central Africa" in place of "Middle Africa".

#### Appendix S2 – Occurrence data

Table S2.1: Pre-processing procedure

| **Table S2.1: Pre-processing of GBIF data** | | |
|---|---|---|
| **Action** | **Number of entries changed** | **Entries remaining** |
| Original GBIF dataset (GBIF, 2023) | | 4270 |
| Filtered to include only entries pertaining to species-level identifications or lower | 566 removed | 3704 |
| Discard entries with no coordinate information, those with identical longitude and latitude, and those with both longitude and latitude equal to zero | 535 removed | 3169 |
| Filtered to include only entries whose coordinates have one or more decimal places | 39 removed | 3130 |
| Discard entries that fall exactly on a country centroid, in GBIF headquarters, and biodiversity institutions | Country centroid: 49 removed<br>(GBIF HQ: 0)<br>(Biodiversity institutions: 0) | 3081 |
| Temporarily filter entries which fall in countries other than the one listed in the record or in the sea | 123 removed | 2958 |
| From the 123 removed entries, filter those that fall in the sea. If these are within a buffer of 0.1 degree (approximately 10km) from land belonging to the country in the record, assign entry to the closest land and replace in dataset | (In the sea: 12)<br>(Within buffer: 11)<br>In correct country when assigned: 10 replaced | 2968 |
| Manual check of remaining records which fell in countries other than that in the record. Those with correctable errors rectified and replaced in the set (excluding those in Asia) | 64 replaced<br>(Asian species not included in this step) | 3032 |
| Crop to project extent | 1333 | **1699** |
| Note: Nineteen entries were removed from the revision dataset when cropped to the project extent, resulting in a final total of 4979 | | |

Table S2.2: Taxonomic standardisation of GBIF data

**Table S2.2: Taxonomic standardisation of GBIF data according to H. Takano (African Natural History Research Trust, personal communication)**

| Action | Number of entries changed | Reason |
|---|---|---|
| *C. birmanensis* removed | 11 | Not a recognised African species |
| *C. fastidiosus* removed | 2 | Placed into *incertae sedis* |
| *C. ninus* to *C. approximans* | 1 | *C.ninus* newly synonymised with *C. approximans* |
| *C. oedipus* to *C. polynices* | 12 | *C. oedipus* newly synonymised with *C. polynices* |
| *C. platycerus* to *C. obtusicornis* | 23 | *C. platycerus* designated as a junior primary homonym |
| *C. simillimus* to *C. polynices* | 5 | *C. simillimus* newly synonymised with *C. polynices* |
| *C. vansoni* to *C. longiceps* | 1 | *C. vansoni* newly synonymised with *C. longiceps* |
| *C. philus* to *C. vitulus* | 125 | *C. vitulus* newly synonymised *C. philus* |
| *C. pseudooedipus* to *C. princeps* | 1 | *C. pseudooedipus* newly synonymised to *C. princeps* |
| **After these changes, 1686 entries remained in the GBIF set** | | |
| Note: no records were re-identified as part of this process, and only the species names of these records were changed. Only records whose original names have been subject to revision were altered in this way, and these changes do not encompass all alterations made in the revision. Taxonomic changes as a result of this revision are pre-publication. | | |

Table S2.3: Manual identification and removal of duplicates in creation of the combined set

When the combined set was created, duplicate records were manually identified and removed. Criteria used to identify duplicates are as follows:

| **Table S2.3:** Manual identification and removal of duplicates in creation of the combined set | |
|---|---|
| **Criterion** | **Explanation** |
| Species name and year identical | If year = NA, this was automatically deemed not a duplicate. Many collecting locations were returned to year after year, and so, without this information, comparison between records was impossible |
| Lat and long <1 degree apart | Could not ensure that duplicates had completely identical coordinates given differing levels of precision used by collectors and institutions when records were digitised and / or uploaded to GBIF |
| If date listed with more precision than just year, these must be identical | |
| Collector and location the same, even if written in a different way | When digitised and / or uploaded to GBIF, label information is formatted in a way that may be different from the original specimen label. As the revision dataset uses original label information, the format of the contents may be different, but the content must be the same to be considered a duplicate |
| No more than two columns can differ in any way | Although more columns were included in each separate dataset, those that were comparable contained information on: Species, year, recorder / collector, longitude, latitude, collecting location (description), date and total number of specimens<br>Some records fulfilled all criteria for duplicates with the exception of total number of specimens. In these cases, if the revision dataset total was higher, the original revision dataset entry was removed and the difference in total re-entered as a new record.<br>E.g. Revision entry with a total of 5 specimens fulfils all the criteria for being a duplicate of a GBIF entry with total of 3 specimens. The original revision entry is removed, but an identical record created with a total of 2 specimens, to ensure all extra value from the taxonomic revision is included.<br>This was thought to be likely a consequence of the digitisation and mobilisation process, in which specimens were missed or even added to collections after this had taken place |

**Appendix S3 – Environmental data**

Table S3.1: WorldClim Variables used in the principal components analysis (Fick and Hijmans, 2017; WorldClim, 2020)

| **Table S3.1**: WorldClim Variables used in the principal components analysis (Fick and Hijmans, 2017; WorldClim, 2020) | |
|---|---|
| **Code** | **Description** |
| BIO1 | Annual Mean Temperature |
| BIO2 | Mean Diurnal Range (Mean of monthly (max temp - min temp)) |
| BIO3 | Isothermality (BIO2/BIO7) (×100) |
| BIO4 | Temperature Seasonality (standard deviation ×100) |
| BIO5 | Max Temperature of Warmest Month |
| BIO6 | Min Temperature of Coldest Month |
| BIO7 | Temperature Annual Range (BIO5-BIO6) |
| BIO8 | Mean Temperature of Wettest Quarter |
| BIO9 | Mean Temperature of Driest Quarter |
| BIO10 | Mean Temperature of Warmest Quarter |
| BIO11 | Mean Temperature of Coldest Quarter |
| BIO12 | Annual Precipitation |
| BIO13 | Precipitation of Wettest Month |
| BIO14 | Precipitation of Driest Month |
| BIO15 | Precipitation Seasonality (Coefficient of Variation) |
| BIO16 | Precipitation of Wettest Quarter |
| BIO17 | Precipitation of Driest Quarter |
| BIO18 | Precipitation of Warmest Quarter |
| BIO19 | Precipitation of Coldest Quarter |

Figure S3.1: Distinct climate types identified in the study area by converting PCA values into classes

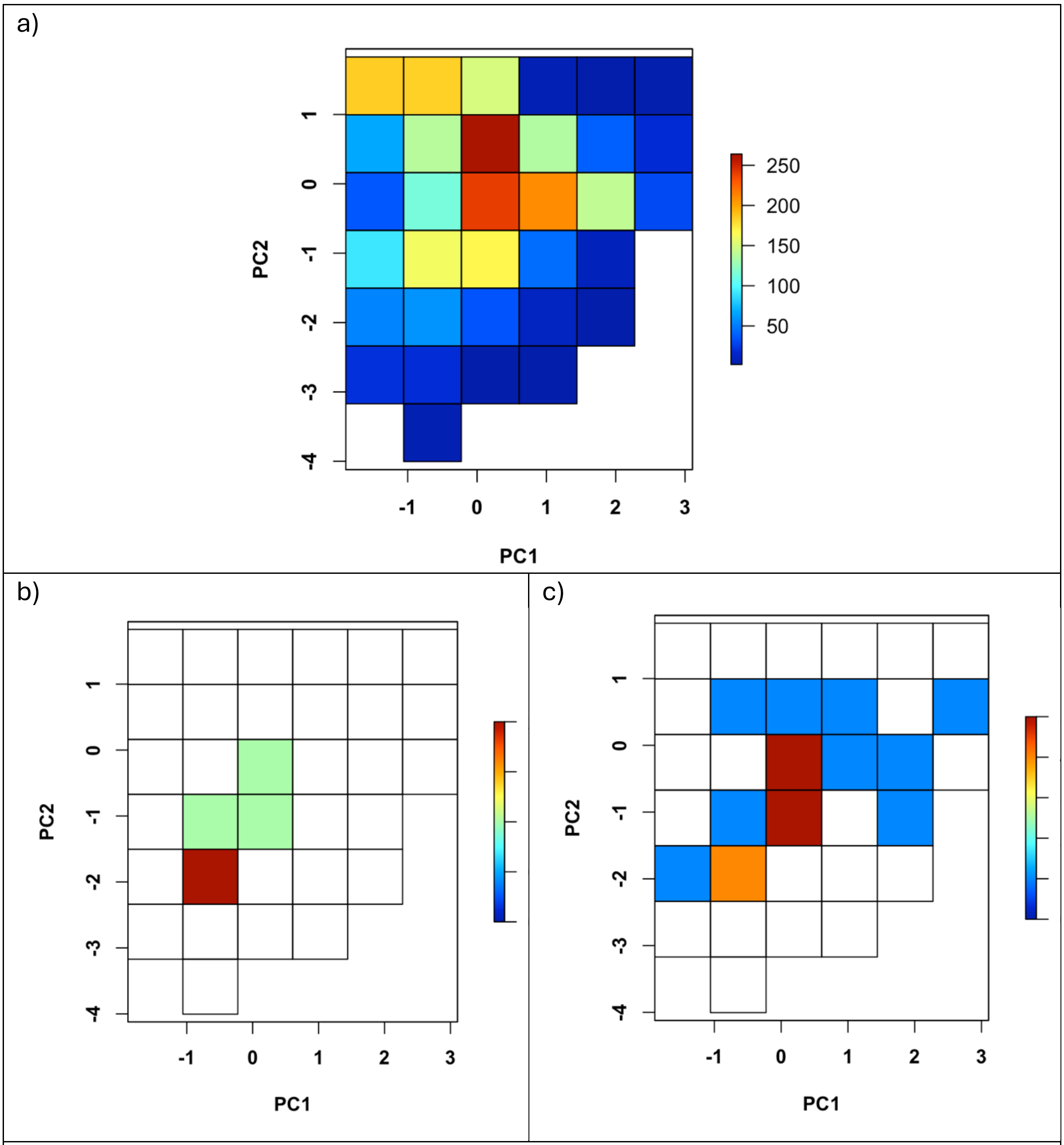

Figure S3.1. Thirty-three climate types were identified in the Afrotropical mainland when principal components analysis values were converted into classes (a). Climate types found in well-sampled *Catharsius* cells for the GBIF set (b) and combined set (c) are displayed with colour indicating the number of cells that particular climate type was found in.
